# Imaging Renal Ultrastructure using a Fast and Simple Optical Clearing and Swelling Protocol

**DOI:** 10.1101/2020.07.10.196733

**Authors:** David Unnersjö-Jess, Linus Butt, Martin Höhne, Anna Witasp, Lucas Kühne, Peter F. Hoyer, Jaakko Patrakka, Paul T. Brinkkötter, Annika Wernerson, Bernhard Schermer, Thomas Benzing, Lena Scott, Hjalmar Brismar, Hans Blom

**Affiliations:** Science for Life Laboratory, Dept. of Applied Physics, Royal Institute of Technology, Solna, Sweden; Department II of Internal Medicine and Center for Molecular Medicine Cologne (CMMC), University of Cologne, Germany; Cologne Excellence Cluster on Cellular Stress Responses in Aging-Associated Diseases (CECAD), University of Cologne, Germany; Division of Renal Medicine, Department of Clinical Sciences, Intervention and Technology, Karolinska Institutet, Stockholm, Sweden and Clinical Pathology and Cytology, Karolinska Universiy Hospital Huddinge, Stockholm Sweden; Pediatric Nephrology, Pediatrics II, University of Duisburg-Essen, Essen, Germany; KI/AZ Integrated CardioMetabolic Center, Department of Laboratory Medicine. Karolinska Institutet at Karolinska University Hospital Huddinge, Stockholm, Sweden; Science for Life Laboratory, Dept. of Women’s and Children’s Health, Karolinska Institutet, Solna, Sweden

## Abstract

Many light-microscopy protocols have in recent years been published for visualization of ultrastructure in the kidney. These protocols present researchers with new tools to evaluate both foot process anatomy and effacement, as well as protein distributions in foot processes, the slit diaphragm and in the GBM. However, these protocols either involve the application of different complicated super-resolution microscopes or lengthy sample preparation protocols. We here present a fast and simple, 5-hour long procedure for the full three-dimensional visualization of podocyte foot processes using conventional confocal microscopy. The protocol combines and optimizes different optical clearing and tissue expansion concepts to produce a mild swelling, sufficient for resolving foot processes using a diffraction-limited confocal microscope. We further show that the protocol can be used for common visualization of large-scale renal histology, pathology, and kidney protein distributions. Thus, we believe that our fast and simple protocol can be beneficial for routine conventional microscopic evaluation of kidney tissue integrity both in research and in the clinic.

## INTRODUCTION

In recent years, a range of novel optical imaging and sample preparation protocols have been presented both by our group and others, handing kidney researchers and pathologists the opportunity to use light microscopy for the analysis of renal ultrastructure^1–8^, reviewed in^9,10^. The power of these new techniques as tools for quantifying renal ultrastructure is exemplified in recent publications from our group and others^3,11–13^. Importantly, super-resolution STED microscopy together with tissue clearing was a crucial tool for establishing the newly proposed model for albumin filtration referred to as the GBM compression model^11,14^. Although powerful, all of these optical protocols require the use of advanced diffraction-unlimited microscopy techniques, such as stimulated emission depletion (STED) microscopy^5^, single-molecule localization microscopy (SMLM)^4^ or structured illumination microscopy (SIM)^2,3^, or involve the introduction of complex polymer chemistry into the sample to clear and/or physically expand it^1,6,7^. Many of the protocols are also very tedious in duration and require a substantial amount of hands-on labor. This extra layer of complexity might be what is keeping these new methods from being routinely used by both researchers and clinical pathologists. Further, many of the protocols lack access to deep 3D imaging of foot processes (FPs) beyond 5-15 µm in depth^1–3,7,8^. Therefore, we set out to develop a drastically simplified and fast all-optical protocol, which clears and spatially swells kidney tissue samples sufficiently to allow for 3D high resolution analysis of renal anatomy and pathology using a conventional diffraction-limited microscope available in almost all life-science and pathology institutes.

Previous protocol simplification have shown that the introduction of acrylamide polymers into PFA-fixed samples can be totally omitted, while still preserving tissue integrity upon SDS clearing^15^. Further, another publication shows that tissue samples can be expanded more than a factor of two using a simple polymer-free protocol based on immersion in different aqueous solutions^16^. On the basis of this, we removed acrylamide infusion from our more tedious and complex published protocols^5,6^, and changed the sample embedding medium to allow for a mild (1.3-fold linear) swelling of kidney tissue samples. By also optimizing imaging parameters, we achieved high enough effective resolution for three-dimensional visualization of podocyte FPs and the GBM using a conventional confocal microscope.

## RESULTS

### Optical clearing and mild swelling of kidney tissue

Kidney samples were treated according to our simple clearing/swelling protocol, schematically depicted in Fig. 1a. Vibratome sectioning of kidney samples and biopsies is not absolutely necessary, but is helpful for later staining (shorter incubation times), mounting (easier to mount flat samples) and imaging (more sample volume accessible close to objective). It should however be stressed that the agarose embedding and Vibratome sectioning takes maximum 15 minutes for a biopsy-sized sample (∼3-4 slices per full biopsy) and is thus neither a complex nor time-consuming addition to the protocol. If no Vibratome is available, samples can also be cut manually with a razor blade, although it is difficult to cut samples thinner than around 0.5 mm using this method. Of note is that the SDS de-lipidation step was not primarily added for optical transparency, but to ensure sufficient swelling of samples^16^. Further, if omitting the SDS de-lipidation, we show that staining quality becomes poor (Fig. S1a), as has also been shown previously by our group^5^. Moreover, to minimize the time needed for sufficient de-lipidation, the temperature was raised compared to our original optical clearing protocol. De-lipidation of the kidney tissue at 70°C for 1 hour produced good results without sample degradation occurring at even higher temperatures (Fig. S1b). Furthermore, as shown by Murakami et al.^16^ urea and urea-like molecules can effectively swell tissue samples resulting in an increase in effective spatial resolution. 8 M urea embedding indeed resulted in a linear swelling of 1.3 times (data not shown) but was not compatible with the lectin staining we wanted to use for the visualization of both glomerular filtration barrier structures and large-scale histology. We thus lowered the urea concentration to 4 M in a mixture with 80% wt/wt fructose (dubbed FRUIT)^17^, which preserved a linear swelling factor of about 1.3 (Fig. 1b) while still allowing lectin labelling (later see Fig. 2-5). In early versions of the simpler and faster protocol, staining of specific proteins (nephrin/podocin/collagenIV) was carried out using primary and secondary antibodies which had to be incubated 2 hours each to allow for sufficient penetration. To shorten this time, amine-reactive fluorophores were directly conjugated to primary antibodies using NHS chemistry. Initially, using the primary conjugated antibodies in a standard staining buffer (PBST) the signal and labelling efficiency was not high enough for outlining individua foot processes. However, by diluting antibodies in an alternative staining buffer called Scale-CUBIC-1 (10 mM HEPES + 10% Triton-X), which has been reported to increase the signal around 2-fold in immunostaining applications^18^, staining quality was increased to a satisfactory degree (Fig. S1c). Thus, the protocol was further simplified and the duration could be decreased by 2 hours, resulting in a protocol taking only 5 hours from collection of the tissue to acquired image. To compare this protocol to known protocols for visualizing podocyte foot processes, several essential parameters were evaluated (Fig. 1c-f, boldface black font = conventional microscopy protocol, gray font = super-resolution or electron microscopy protocol, our protocol highlighted with a square). Evaluated parameters include number of protocol steps, number of reagents used, hands-on time and total protocol duration. Compared to previously published protocols using conventional microscopy, our fast and simple protocol outperformed them on all evaluated parameters. The protocol presented by Pullman et al.^2^ is slightly faster in total, but with the drawbacks that the protocol requires super-resolution SIM microscopy, more hands-on labor, more sample interactions and number of reagents. Further, the later protocol is performed on thin cryo-sections, thus lacking access to large in situ imaging depths. Of note is that our protocol is less complex than standard paraffin-embedding or cryo-sectioning when all evaluated parameters are considered.

**Figure 1.**
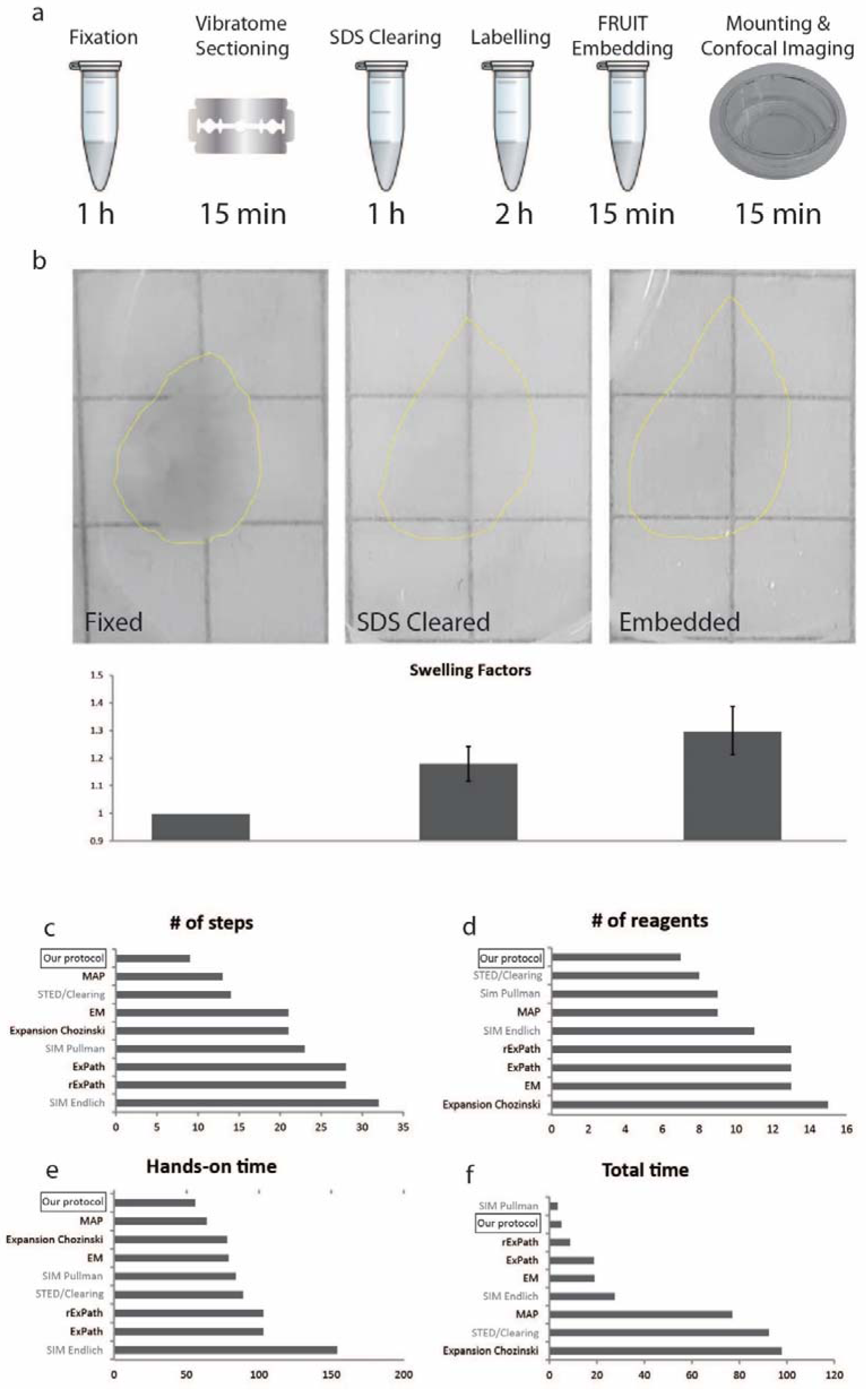
Rapid clearing and mild swelling of kidney samples. (a) General scheme and timeline of our protocol. (b) A 300 µm thick slice of adult rat tissue at different steps of the protocol with corresponding swelling factors. (c-f) Comparison of our protocol to existing protocols^1–3,5–7,31^. Number of steps (c) corresponds to the number of physical interactions with the sample. Number of reagents (d) are the number of reagents needed to perform the protocol. Standard buffers or chemicals, such as PBS are not counted. Hands-on time (e) is the *estimated* time of active work needed to perform the protocol. The total time (f) is the total duration of each protocol.

**Figure 2.**
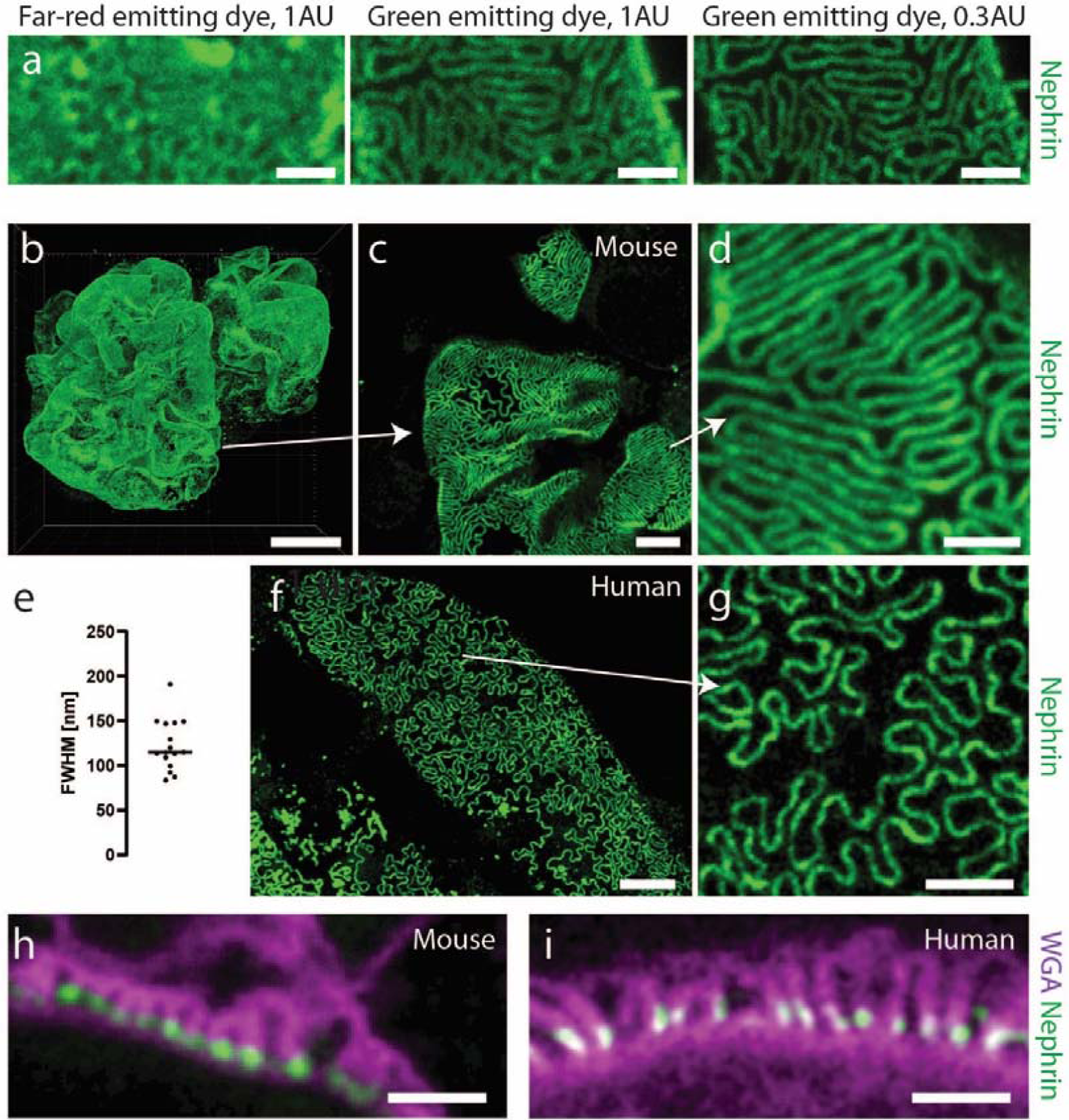
Validation of the protocol for imaging renal ultrastructure. All mouse samples were stained for nephrin using Alexa-488 (a-d, h) or Alexa-405 (f-g,i). Samples on bottom row were additionally stained with WGA lectin conjugated to Alexa-488 (i) or CF-405M (h). All samples were imaged using a Leica SP8 confocal microscope with a 100X/1.4NA oil objective. All images apart from (h-i) are maximum intensity projections of ∼ 2 µm (a, c-d, f-g) or ∼ 40 µm (b) thick z-stacks acquired at depths of 0-50 µm. Scale bars 10 µm (b), 5 µm (f), 2 µm (c, g), 1 µm (a, d, h-i). (a) Images of a mouse kidney sample stained for nephrin using Abberior STAR 635P or Alexa-488, showing the effect on resolution when choosing a dye with shorter excitation/emission wavelength and shutting down the pinhole from 1 to 0.3 airy units. Raw data. (b) Nephrin pattern in a whole glomerulus from a 20 weeks old wild-type mouse, showing expected dense slit diaphragm pattern in glomerular capillaries. (c-d) Zoomed images of the glomerulus in (b) showing that foot processes can be clearly resolved using diffraction-limited confocal microscopy. (e) Effective resolution of the protocol. Line profiles were drawn perpendicular to slit diaphragms, and a Gaussian function was fitted to each nephrin intensity profile. The resolution was estimated as the full-width at half-maximum (FWHM) of the Gaussian fit divided by the swelling factor. (f-g) Slit diaphragm imaging in human kidney. The sample was taken from an adult patient that underwent nephrectomy due to renal carcinoma and imaging was performed far away from the cancer tissue and is thus estimated to correspond to adult healthy kidney tissue. The SD pattern appears looser, which is expected in human kidneys, especially in older patients. (h-i) Cross sectional views of foot processes in the same mouse (h) and human (i) samples as in (b-g). Labelling using wheat germ agglutinin (WGA) shows staining positivity on apical side of podocyte foot processes (glycocalyx) and nephrin in the slit diaphragm.

**Figure 3.**
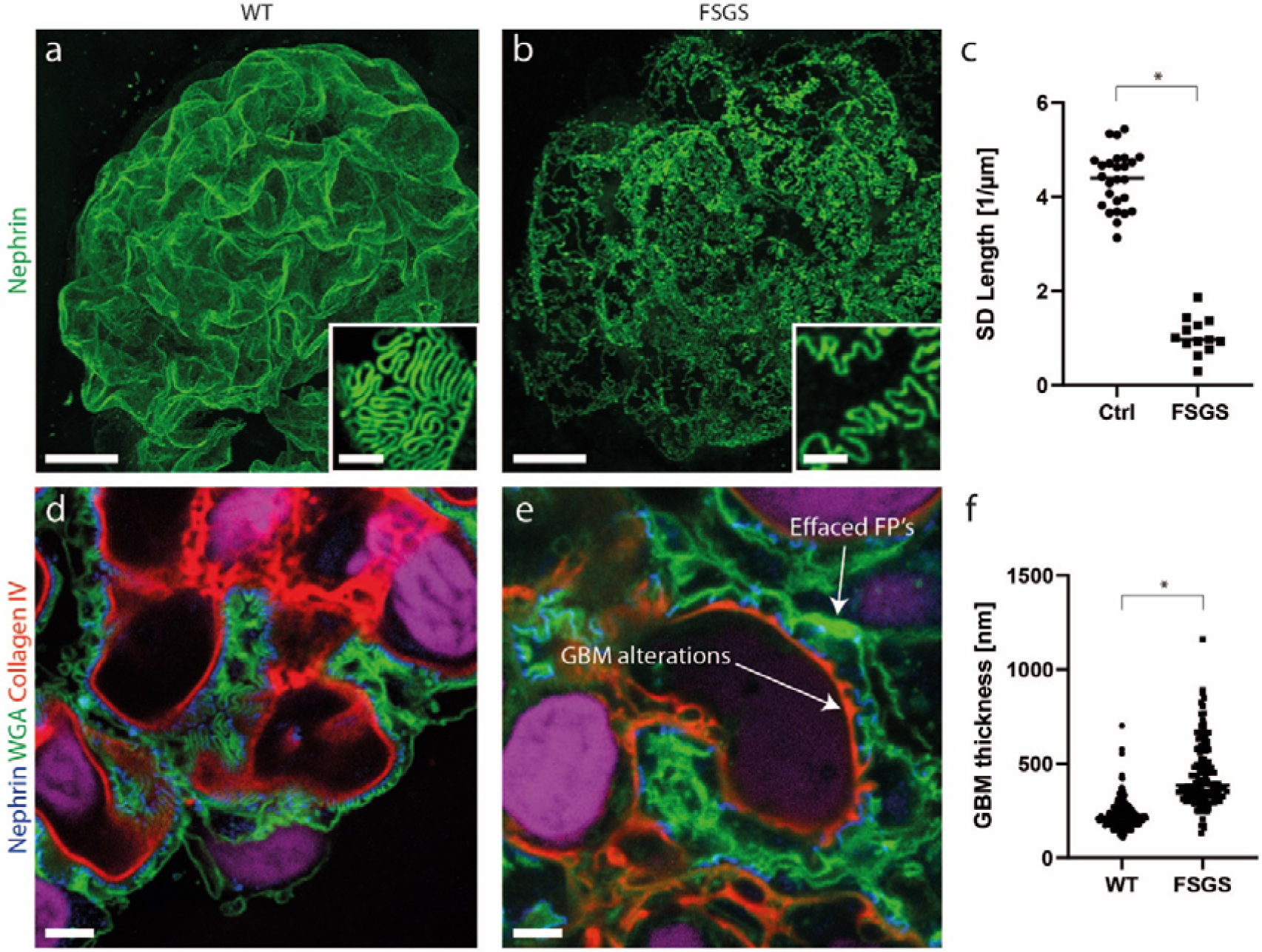
Visualization of renal ultrastructure in a mouse model for FSGS. All samples were stained with an Alexa-488 conjugated nephrin antibody, and Alexa-555 conjugated collagen-IV antibody and CF-405M conjugated WGA. Scale bars 10 µm (a-b, insets 1 µm), 2 µm (d-e). (a-b) Maximum intensity projections of 40 µm thick z-stacks showing the global SD density with an apparent decrease in the FSGS mice. Insets show zooms of glomerular capillaries showing effacement more in detail. (c) SD length per capillary area in WT and FSGS mice, showing an around 4-fold decrease for mutated mice. Two-tailed t-test, P < 0.0001. (d-e) Cross-sectional images of glomerular capillaries showing both effacement (wider foot processes, less nephrin), as well as a thickened, “bumpy” GBM as visualized by the collagen-IV staining. (f) Measurements of GBM thickness in WT and FSGS mice showing both a thickened and more heterogeneous (larger spread of the data) GBM. Two-tailed t-test, P < 0.0001.

**Figure 4.**
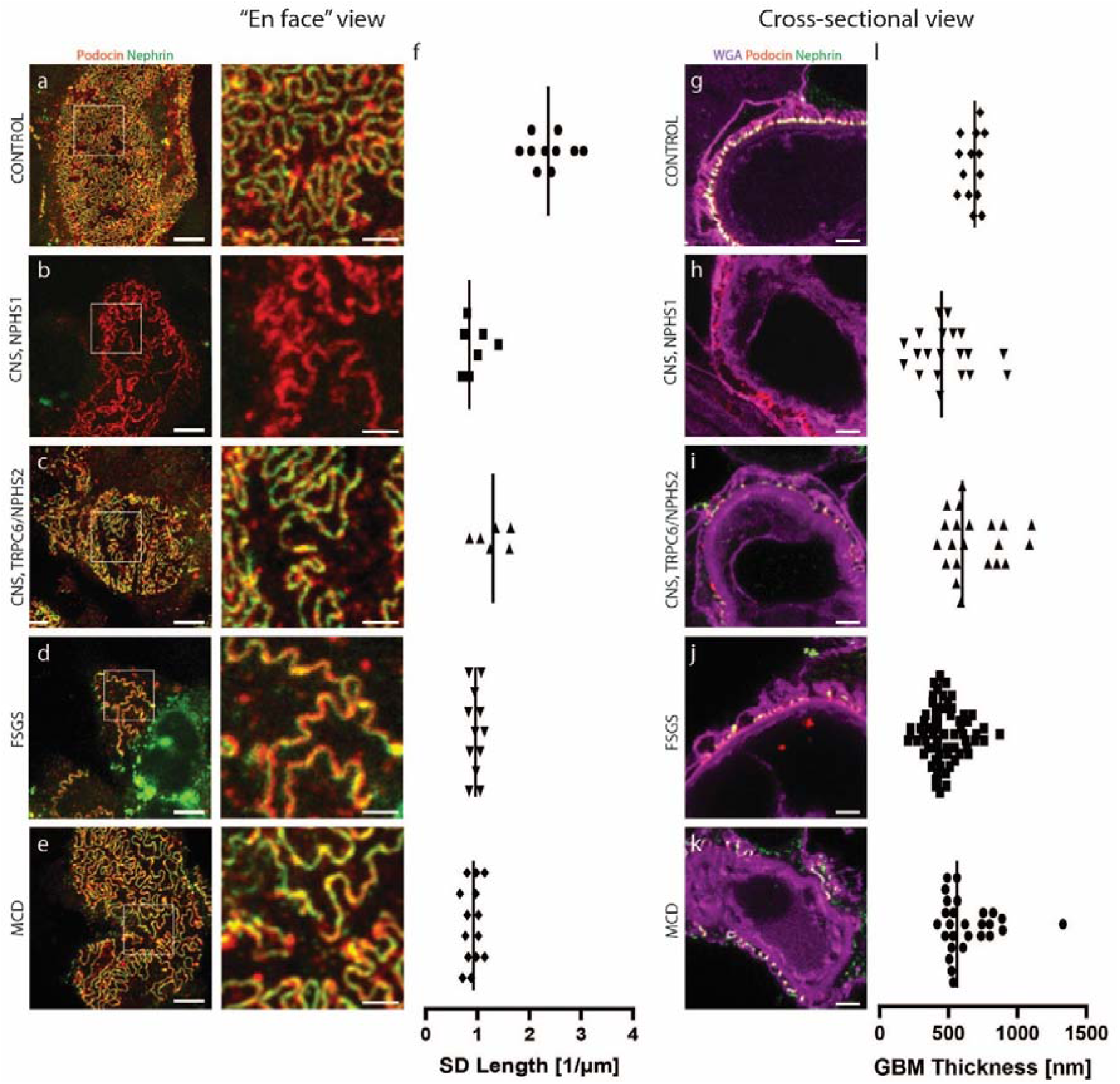
Foot process and filtration barrier pathology in patients with different types of kidney disease. All images acquired using a Leica SP8 confocal system using a 100X NA 1.4 oil objective and a pinhole setting of 0.3 AU. (a-e) Maximum intensity projections of ∼2 µm thick z-stacks in samples from control patient (a), from patients with mutations in the NPHS1 (b) and TRPC6/NPHS2 (c) genes and in patients diagnosed with focal segmental glomerulosclerosis (d) and minimal change disease (e) showing clear foot process effacement in all diseased samples, and expected loss of nephrin for the patient with an NPHS1 mutation. Right column shows zoomed images of the areas indicated with a white square box. Scale bars 3 µm and 1 µm (zoomed images) (f) Slit diaphragm length per area as calculated for the corresponding patients in each row. Measurements from at least 5 different glomerular capillaries in 3 different glomeruli. (g-k) Cross sectional view of the filtration barrier for the same patients as in (a-e) showing foot process effacement in all patients (reflected both by the WGA staining showing wide foot processes and by the decreased amount of SD’s visualized by podocin/nephrin staining). Scale bars 1 µm. (l) GBM thickness measured for the corresponding patients in each row, showing a larger spread of the data for all diagnoses as compared to control, indicating a more heterogeneous and “bumpy” GBM. At least 11 GBM measurements from at least three different glomerular capillaries.

**Figure 5.**
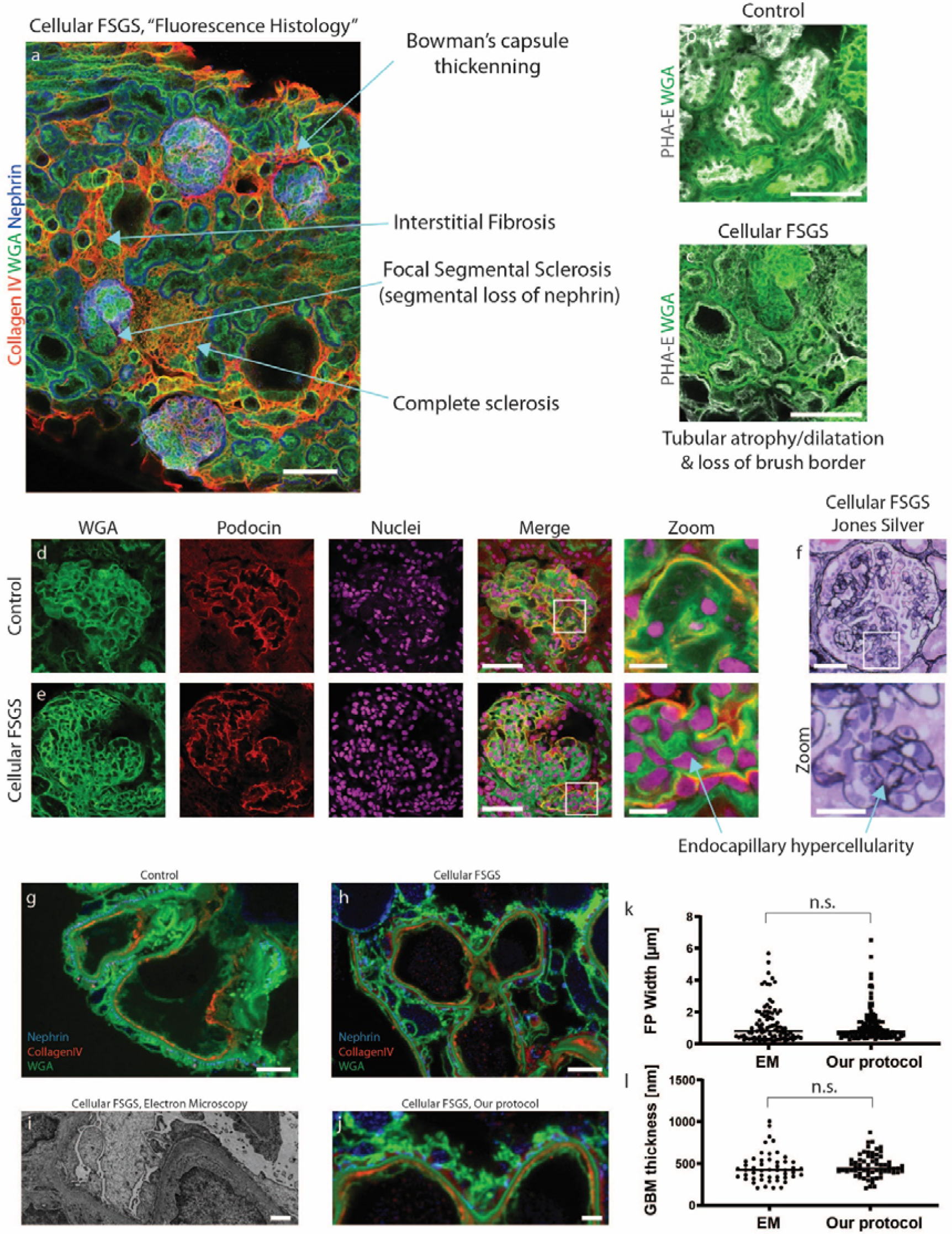
Patient diagnosed with FSGS of the cellular type with comparison to standard pathology methods. All images acquired using a Leica SP8 confocal system. All samples were stained for nuclei with DRAQ-5, nephrin with Dylight-405, glycoconjugates with WGA-Alexa-488, collagen IV or podocin with Cy3 and PHA-E conjugated to Atto-594. Relevant channels are shown as indicated to visualize desired features in each image. (a) Maximum intensity projection of a 15 µm thick confocal z-stack with arrows showing pathological parameters typically seen in FSGS patients are clearly visualized. 10X NA 0.45 air objective. Scale bar 100 µm. (b-c) Confocal images of samples from control (b) and FSGS (b) patients, showing typical tubular pathologies such as dilatation of tubular lumen and also loss of brush border. 10X NA 0.45 air objective. Scale bars 100 µm. (d-e). Confocal images of whole glomeruli from control (b) and FSGS (b) patients clearly showing the endocapillary hypercellularity typical for FSGS of the cellular type. Scale bars 50 µm and 10 µm (zoomed images). (f) Transmitted light microscopy image of glomerulus from the same FSGS patient shown for comparison. Jones Silver stain. Scale bars 50 µm (top), 20 µm (bottom, zoomed). (g-h) Confocal images of samples from control (g) and FSGS (h) patients, showing obvious effacement of foot processes in the FSGS patient, whereas the GBM appears normal. Scale bars 3 µm. (i-j) Comparison between electron microscopy (i) and confocal (j) images of a cross-section of the glomerular filtration barrier from the patient diagnosed with cellular FSGS. Scale bars 1 µm. (k-l) Foot process width (k) (N = 220 (EM), N = 125 (our protocol) measured in at least 3 different capillaries) and GBM thickness (l) (N = 50 (EM), N = 63 (our protocol) measured in at least 3 different capillaries) as measured with our protocol or EM. Two tailed t-tests. p-values 0.44 (k) and 0.51 (l).

### Optimization and validation of our protocol for resolving foot processes

Because our fast and simple protocol gives only a moderate swelling factor of 1.3, confocal imaging parameters had to be optimized to achieve sufficient resolving power for imaging foot processes. Mouse kidney samples were treated according to the protocol and stained for nephrin using different fluorophores. As is common knowledge, the resolution of a confocal microscope depends both on the pinhole size and the wavelength of the excitation/emission light^19^. We thus used a smaller pinhole diameter of 0.3 airy units (AU) together with a green-emitting dye Alexa 488, and in this way achieved high enough resolution to resolve foot processes in a wild-type mouse kidney (Fig. 2a). Thus, whenever mouse foot processes are to be visualized, fluorophores with an excitation wavelength of maximum 488 nm (e.g. Alexa-488, Alexa-405, CF405M) are used. With these optimized imaging conditions, we were able to visualize whole mouse glomeruli in 3D, at an effective resolution of around 115 nm, well enough to resolve individual foot processes (Fig. 2b-e). Moreover, we show that with the two-hour long primary labelled antibody incubation step, a penetration depth of at least 70 µm is achieved (Fig. S2a-c). Using a glycerol objective with a correction collar, this gives access to larger z-depths than any of the previously developed protocols, with the exception being our clearing/STED protocol^5^. Since the resolution of a confocal microscope is highest in the xy-dimension, capillaries must preferentially be oriented in this imaging plane for optimally resolving foot processes. This imaging condition holds true for all the other optical protocols mentioned in this article as well. Keeping the glomerular capillaries in their 3D *in situ* setting allows us to screen through the whole glomerulus. We show that in every glomerulus, there are around 0.2 (human) to 0.3 (mouse) xy-oriented areas per µm in z (Fig. S2d). This implies that we can with our protocol find plenty (around 10 large field-of-view areas), allowing us to quantify and easily follow also subtle variations in renal nanoscale structures^11,14^. This shows the importance of having access to the depth-dimension when performing high-throughput quantitative analyses of glomerular health. Importantly, we also validated the protocol for also resolving foot processes in human patient material (Fig. 2e), which is less challenging as compared to mouse, due to the larger dimensions of human FP’s^11,13^. Using wheat germ agglutinin (WGA) lectin with affinity towards the glycocalyx of FP’s, we were moreover able to resolve foot processes also in a cross-sectional “transmission-EM-like” manner, in both mouse and human samples (Fig. 2f-g). Additionally, we co-stained for nephrin, with the expected localization in the slit membrane, showing that we can get data on protein localization in addition to morphological information. Taken together, we have demonstrated that our simple clearing and swelling protocol can in just a few hours provide three-dimensional information regarding renal ultrastructure and protein expression at a resolution previously only accessible with more complex protocols.

### Ultrastructure Pathology in Mouse Samples

As previously shown^2,3,5,6,11,12^, “en face” foot process imaging of the SD can readily be utilized to visualize and quantify foot process effacement. We validate that this rationale can be applied also with our new protocol by using a recently published mouse model for FSGS, with two compound heterozygous point mutations in the podocin gene^11^ (Fig. 3). By staining for nephrin, the SD pattern could be visualized in three dimensions in whole glomeruli of WT and mutated mice (Fig. 3a-b). As previously shown^3,11^, the SD length per area can be semi-automatically quantified using ImageJ macros, and applying this morphometric analysis gave a result consistent with what has previously been published for this mouse line^11^ (Fig. 3c). Further, glomerular capillaries were imaged in cross-section showing both effacement of foot processes as well as alterations to the GBM in the mouse model for FSGS (Fig. 3d-e). The GBM thickness was semi-automatically measured, showing a significant increase in mutated mice modelling FSGS disease progression (Fig. 3f).

### Ultrastructure Pathology in Human Samples

To further validate our protocol for possible clinical use, we applied it to human tissue from patients diagnosed with different types of kidney disease. Two patients with congenital nephrotic syndrome (CNS), either with a homozygous mutation in the NPHS1 gene (encoding for nephrin), or trans-associated mutations in the TRPC6 and NPHS2 genes (encoding for the TRPC6 channel and podocin). We also applied the protocol to two patients diagnosed with focal segmental glomerulosclerosis (FSGS) and minimal change disease (MCD). Using our protocol, we could indeed visualize and quantify typical hallmarks of kidney disease such as foot process effacement as well as pathologies in the GBM (Fig. 4). Images of renal ultrastructure could be acquired both “en face” (Fig 4a-e) and from the side (Fig. 4g-k). We could moreover observe the expected absence of nephrin in slit diaphragms of the patient with a homozygous mutation in the NPHS1 gene^20^ (Fig. 4b,h). As for mouse samples (Fig. 3), filtration barrier morphology could be quantitatively evaluated by measuring the slit diaphragm length per area and the GBM thickness using semi-automatic ImageJ macros (Fig. 4(f, l), S3(a-b)). Of note, expected GBM thickening is not observed in any of the diagnosed patients as compared to control (as observed for mice, Fig. 3f). The explanation likely lies in the fact that GBM thickness increases with age^21^. The nephrectomized control kidney was taken from an older patient while all the other samples are from infants or young children, so GBM thickening would probably be observed if compared to age-match control samples.

### Large-Scale Pathology in Mouse and Human Samples

Since not only ultrastructural, but also histological changes to renal morphology is of high interest both in research and in the clinic, we investigated the possibility of our protocol to assess large-scale morphology using fluorescence-based histological analysis. We show that using a basic 10X/0.45NA air objective, we could detect large scale pathologies such as focal partial or total loss of nephrin (Fig. S4a), global loss of nephrin in glomeruli from the patient with an NPHS1 mutation (Fig. S4a), different degrees of interstitial fibrosis (Fig. S4b), glomerulosclerosis (Fig. S4c), focal thickening of Bowman’s capsule (Fig. S4c), atherosclerosis (Fig. S4d), loss of brush border in tubules (Fig. S4e) and dilatation of tubular lumen (Fig. S4d). Also in the mouse model for FSGS, we were able to detect both segmental sclerosis of glomeruli as well as tubular alterations (Fig S5).

### Visualizing Large- and Ultra-Scale Alterations in Patients Diagnosed with Focal Segmental Glomerulosclerosis (FSGS) and IgA nephropathy

Moreover, to further exemplify the possible use of the protocol in patient samples, we investigated if we could detect pathology features important for diagnosing a case of focal segmental glomerulosclerosis (FSGS) of the cellular type (Fig. 5). We show the visualization of most pathology features mentioned in the clinical scoring statement from the pathologist, such as focal total/segmental sclerosis, tubular atrophy and endothelial hypercellularity in glomerular capillaries when comparing to present gold standard methods such as optical histology and electron microscopy (Fig. 5a-j). Both qualitatively and quantitatively (Fig. 5k-l, S3) we see that our fast and simple protocol provides comparable diagnostic information as standard pathology methods presently used for renal pathology. To further investigate the diagnostic possibilities of our protocol, we also applied it to a biopsy from a patient diagnosed with IgA nephropathy (IgAN) (Fig. 6) and were able to confirm the visualization of many diagnostic features. The biopsy showed no severe pathologies in the filtration barrier as shown both by EM and our protocol (Fig. 6a-b) and a comparative analysis resulted in no obvious differences between EM and our protocol (Fig. 6c-d). Next, the histology of the IgAN kidney was evaluated both with standard histology (Fig. 6e) and “fluorescence histology” (Fig. 6f). We could visualize the alterations (or lack of alterations) mentioned in the statement from the pathologist such as interstitial infiltration of cells (Fig. 6g-h), normal glomerular capillaries (Fig. 6i) and mesangial expansion and hypercellularity (Fig. 6j).Visualization of immune deposits is critical for diagnosing IgAN and other immune complex diseases. Therefore, we stained the IgAN biopsy using an IgA antibody and were able to clearly localize the deposits both on the large scale (Fig. 6k-l) and in detail (Fig. 6m). The staining pattern was equivalent to that of standard immunofluorescence (IF) (Fig. 6n-o). By imaging foot processes “en face”, partial mild effacement could also be visualized (Fig. 6p) and quantified (Fig. 6q). Even though we only got access these two patients where comparisons to standard pathology was available, the results indicate that parallel evaluations using our protocol and standard pathology yield similar pathological data regarding immune deposits (IF), histology and ultrastructure (EM).

**Figure 6.**
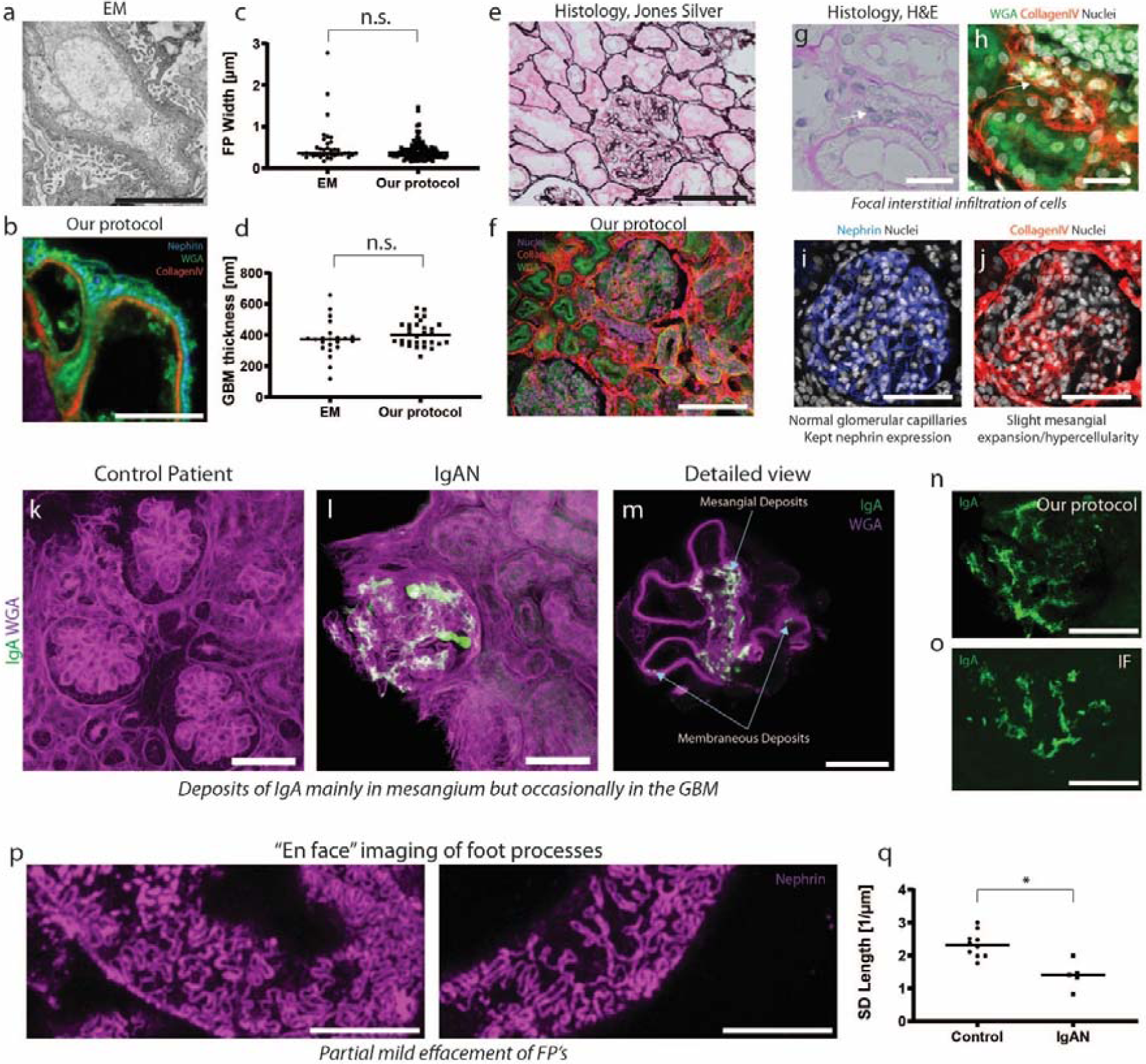
Patient diagnosed with IgAN. 20X NA 0.75 oil (f, h-l, n) or 100X NA 1.4 oil (b, m, p) objectives. Samples were stained either for nuclei (DRAQ-5), nephrin (Alexa-594), glycoconjugates (WGA-Alexa-488) and collagen IV (Cy3) or IgA (FITC) and glycoconjugates (WGA-Alexa647). (a-b) The GFB as visualized by EM (a) and our protocol (b), showing no apparent abnormalities. Scale bars 5 µm. (c-d) GBM thickness (c) (N = 22 (EM), N = 30 (our protocol) measured in 1 capillary for EM and 3 capillaries for our protocol) and FP width (d) (N = 38 (EM), N = 115 (our protocol) measured in 1 capillary for EM and 3 capillaries for our protocol) measurements with EM and our protocol. Two tailed t-tests. p-values 0.24 (c) and 0.16 (d). (e-f) Standard histology (e) and our protocol (f) in the IgAN patient. Image acquired with our protocol shows a maximum intensity projection (MIP) of a 20 µm thick z-stack between 5-25 µm z-depth. Scale bars 100 µm. (g-h) Hematoxylin & eosin stain (g) and our protocol (h) showing interstitial infiltration of cells (arrowheads). Scale bars 20 µm. (i) Whole glomerulus acquired with our protocol showing normal capillaries and nephrin expression. Scale bar 50 µm. (j) Whole glomerulus acquired with our protocol showing mesangial expansion (collagen IV staining) and hypercellularity. Scale bar 50 µm. Images shown in (h-j) taken from the same projection as in (f). (k-l) MIP of 50 (k) and 20 (l) µm thick z-stacks acquired with our protocol of samples from control (k) and IgAN (l) showing strong positivity for IgA in the IgAN patient. Scale bars 50 µm. (m) Image of the same sample as in (l) showing in detail the mesangial (and rare membraneous) deposits of IgA. Scale bar 5 µm. (n-o) Glomeruli stained for IgA using our protocol (n) or standard IF (o) showing equivalent staining pattern. Image acquired with our protocol shows a MIP of a 10 µm thick z-stack acquired at 10-20 µm depth. Scale bars 50 µm. (p) MIP of 2 µm thick z-stacks acquired at 5-10 µm z-depth showing mild effacement of FP’s. (q) Box plots of SD length per area showing a decrease for the IgAN patient. Two-tailed t-test. p = 0.004.

## DISCUSSION

We here present a fast, optimized and simplified way of performing high-throughput visualization of both large-scale (e.g. tubules, glomeruli) and nanoscale (e.g. foot processes, filtration barrier) renal structures. This renal analysis could be achieved in just five hours (from dissection to analysis) which is substantially shorter than most previous clearing and expansion protocols. As a comparison, the turn-around time for full renal diagnostics (LM, IF and EM analysis) is a day or two^22^. In terms of total duration, complexity, hands-on work and number of reagents, our workflow outperforms previously published microscopy protocols for foot process analysis and glomerular filtration barrier imaging in the kidney. Moreover, we show that the resolution we achieve is *high enough* to draw conclusions about pathologies in the filtration barrier using simple confocal microscopy, a standard tool already available in most research facilities and pathology labs. Since the swelling of the samples is mild, the effective field-of-view is larger than for other more tedious expansion protocols, allowing also for high-throughput imaging of large-scale morphology and pathology of human samples. It should be stressed that SDS clearing (de-lipidation) is in the protocol not added primarily for optical transparency, but rather to allow for sufficient labelling and swelling of samples. We show, however, that the optical transparency is beneficial when performing imaging of foot processes as a large imaging volume is accessible, allowing for finding and screening more suitable regions of the sample for quantitative analysis.

Furthermore, an important finding of our study is that our protocol performs complementary and beneficially vs. IF/Cryosections, histology and EM, i.e. protocols presently used for standard analysis of sectioned kidney tissue. Still, we believe that the biggest advantage over standard protocols is that this protocol can potentially merge all three workflows of standard microscopy into just one with apparent savings in terms of working hours and material/reagents/equipment needed.

Recent studies have applied Airyscan microscopy, which utilize variable pinhole settings on a small camera array detector in combination with deconvolution and signal reassignment image processing^23,24^. In this study we show that with mild swelling and subsequent superb staining and imaging conditions, just conventional confocal imaging with a closed-down pinhole provides sufficient resolution without mathematical post-processing of images. Thus, our simple and fast 3D-compatible protocol can be performed with any standard confocal microscope equipped with a suitable high NA objective (oil/NA 1.4 and glycerol/NA 1.3 tested in this work).

For possible use in clinical pathology, the protocol must be further validated for evaluation of the plethora of pathological parameters present in other types of kidney disease as well. As one example, it is in some cases crucial to stain for amyloid deposits (presently visualized using Congo red histology stain) to diagnose amyloid nephropathy. We believe such detection issues are mostly a matter of developing new non-histological staining strategies, and in fact fluorescence-based methodologies for amyloid visualization have been demonstrated by others^25^. Moreover, the resolution of our protocol is not optically sufficient to visualize nanoscale fibrillary deposits, so for some special cases one might still have to apply EM, or a combination of our protocol with super-resolution fluorescence microscopy for accurate diagnosis. In this work we present comparative data between our protocol and EM for the evaluation of glomerular filtration barrier morphology which indicates that the two approaches yield very similar results. However, it should be stressed that the analysis is carried out on a small number of patients, so a more thorough analysis is in the future necessary to further establish this point. To achieve this, a focused collaborative effort involving both pathologists, sample preparation experts and imaging experts has to be initiated. After such further validations and optimizations, a fast and simple all-scale optical analysis might be at hands for use in renal diagnostics. This thorough validation has not been within the scope of this proof-of-principle study, where the aim was to develop a fast optical protocol for kidney ultrastructure imaging.

Presently, development of improved optical kidney diagnostics is explored by several researchers. There have recently been demonstrations that pathological parameters can be extracted from cleared kidney tissue using both non-labelled (autofluorescence) and fluorescently labelled samples^26,27^. Furthermore, a recent study shows that “virtual histology” can be performed using deep learning algorithms to auto-fluorescence images^28^ and neural network-based segmentation and classification of kidney pathologies has also been shown^29,30^. Merging novel sample-preparation methods (as the ones shown in this study) with new image analysis tools poses many interesting digital possibilities in the future, and we could likely see changes to how we image and analyze kidney samples in the clinic soon.

## MATERIALS AND METHODS

### Mouse, rat and human kidney tissue

Rats used in the experiments were purchased from Charles River, Germany. Animals were anesthetized (intraperitoneal injection of pentobarbital), the aorta cut, and kidneys dissected. All experiments were approved by Stockholm North Ethical Evaluation Board for Animal Research and were performed in accordance with animal welfare guidelines set forth by Karolinska Institutet.

All mouse experiments were approved by the State Office of North Rhine-Westphalia, Department of Nature, Environment and Consumer Protection (LANUV NRW, Germany) and were performed in accordance with European, national and institutional guidelines. Mice of 100% C57BL/6N background were used. After anesthesia with Ketamine and Xylazine, mice were euthanized by cardiac perfusion with Hank’s Balanced Salt Solution (HBSS) and fixated as described below.

Patient material was obtained from two children suffering from Steroid-Resistant Nephrotic Syndromes due to either point mutations in NPHS1 or compound-heterozygous point mutations in TRPC6 and NPHS2, as well as three patients diagnosed with focal segmental glomerulosclerosis, minimal change disease and IgA nephropathy, respectively. Control human tissue was collected from patients that were nephrectomized due to renal tumors. Tissue sample was dissected from the non-tumorous pole of the kidney and showed normal histological picture in routine histological examination. All procedures were approved by the Ethics Commission of Cologne and the regional Ethical Committee of Stockholm and conducted in accordance with the declaration of Helsinki. When applicable, patients or the patient’s parents gave informed consent.

### Mouse Model for FSGS

Mice with two compound-heterozygous point mutations, Pod^R231Q/A286V^, were generated as previously described^11^. Mice were sacrificed at 20 weeks of age as stated above.

### Fixation

Human, rat and mouse kidneys were fixed in 4% PFA in 1X PBS at room temperature for 1-3 hours. The human kidney tissue from a renal carcinoma patient was fixed in Duboscq-Brasil solution for 3 hours, stored in 70% EtOH and hydrated in 1X PBS for 5 minutes prior to the protocol.

### Sectioning

If needed, samples were embedded in 3% agarose and sectioned using a Vibratome to 300 µm thickness.

### De-lipidation

Lipids were removed from kidney samples by incubation in clearing solution (200 mM boric acid, 4% SDS, pH 8.5) at 70°C for 1 hour with shaking at 500 rpm.

### Labelling

Samples were washed for 5 min in 1X PBS with 0.1% TritonX-100 (PBST) prior to immunolabelling to remove SDS. Samples were then incubated in primary antibody (+ lectins, DRAQ5 for latest version of protocol (primary conjugated antibodies)) diluted in PBST (older version) or 10nm HEPES with 10% TritonX-100 (Scale-CUBIC-1, newer version) at 37°C for 2 hours with shaking at 500 rpm, followed by washing in PBST for 5 minutes at 37°C. For the latest version of the protocol with primary conjugated affinity molecules, the staining protocol was terminated here. For the earlier version (primary + secondary) samples were then incubated in secondary antibody (if primary AB not conjugated to fluorescent dye), lectins and DRAQ-5 diluted in PBST at 37°C for 2 hours with shaking at 500 rpm, followed by 5 minutes washing in PBST at 37°C.

### Mounting

Samples were incubated in 75% wt/wt fructose (1 mL of de-ionized water added to 4g of fructose) containing 4M Urea at 37°C with shaking at 500 rpm for 15 minutes and then placed in a MatTek dish with a cover slip on top (to prevent evaporation) prior to imaging.

### Antibodies and probes

DRAQ-5 (Thermo Fisher, 62251) was used at a dilution of 1:500. Wheat Germ Agglutinin (Thermo Fisher, W11261) and PHA-E (Vector Laboratories, B-1125) was conjugated to Alexa-488 (Thermo Fisher, A20000) and Atto-594 (AttoTech, AD 594-31), respectively, using NHS chemistry and were used at a 50 µg/mL concentration. Also, WGA conjugated to CF405M (Biotium, 29028-1) was used at the same concentration. Primary antibodies used in this study were rabbit polyclonal antibody to podocin (Sigma-Aldrich, P0372) used at a dilution of 1:100, sheep polyclonal antibody to nephrin (R&D systems, AF4269) used at a dilution of 1:50, rabbit polyclonal antibody to collagen IV (Abcam, ab6586) used at a dilution of 1:50 and a FITC-conjugated rabbit polyclonal antibody to IgA (Dako, F0204). The polyclonal sheep anti-nephrin antibody was also directly conjugated to Alexa-488 using NHS chemistry. A rabbit monoclonal antibody to collagen IV (Abcam, ab256353) was conjugated to Alexa-555 using NHS chemistry. Secondary antibodies used in this study were donkey anti-goat conjugated to Alexa-405 (Abcam, ab175664), DyLight-405 (Jackson Immuno Research, 705-475-003), Alexa-488 (Thermo Fisher, A-11055), Alexa-594 (Thermo Fisher, A-11058) or Abberior STAR635P (Abberior, 2-0142-007-2, dilution 1:50) and donkey anti-rabbit conjugated to Alexa-555 (Thermo Fisher, A-31572) or Cy3 (Jackson Immuno Research, 711-165-152), all used at a dilution of 1:100 unless other stated.

### Imaging

Images were acquired using a Leica SP8 confocal system (20X NA 0.75 multi-immersion objective (oil used) or 100X NA 1.4 oil objective as stated in figure legends). Imaging with the 100X objective always implies a pinhole setting of 0.3 AU (as calculated for 594 nm light) if not stated otherwise. Standard histology imaging was performed using a Leica DM3000 system (Fig. 5) or an Olympus BX45 system (Fig. 6), standard immunofluorescence was performed using a Nikon Microphot-FXA and EM performed with a Zeiss 912 (Fig. 5) or a Hittachi 7700 (Fig. 6) transmission electron microscope.

### Image processing and analysis

Images were smoothened (unless other stated) by replacing each pixel value by the mean of its 3×3 neighborhood using Fiji/ImageJ. The morphometric analyses described below are visually presented in Supplementary Figure S3.

### Slit diaphragm length measurement

Slit diaphragm (SD) length per area was determined using ImageJ/Fiji. In brief, the fluorescent signal was enhanced using background subtraction, mean filtering and contrast enhancement. The SD signal was detected with the help of the ridge detection plugin as previously published by us and Siegerist^3,5,11^. In the IgAN-patient (Fig 5 p,q), the measurement of slit diaphragm length was done manually. This macro can be downloaded from GitHub at https://gist.github.com/github-martin/e699d18ae6cfa5b1a6aace15d3c3544c.

### GBM thickness measurement

The GBM thickness was evaluated using an ImageJ macro where the inner and outer boundaries of the GBM were manually assigned based on the collagen IV and nephrin staining or for EM images based on the electron density grayscale (see Fig. S3).

### Slit diaphragm width

For EM images slit diaphragms were visually identified and the distance between them were manually measured using line profiles in ImageJ. The slit diaphragms were identified as nephrin-positive “spots” along the filtration barrier and the distance between them were measured in the same way as for EM images.

### Schemes

The scheme in Fig. 1a was produced by D.U-J apart from the vector graphics for the Eppendorf tube and the razor blade which were downloaded from https://pixabay.com/vectors/vial-tube-fluid-laboratory-41375/ and https://pixabay.com/illustrations/razor-blade-blade-sharp-cutting-220323/ respectively (free to use).

## ACKNOWLEDGEMENTS

This study was supported by grants from the Swedish Research Council (VR 2013-6041) and infrastructure development support from the Swedish Foundation for Strategic Research (RIF14-0091). We thank the CECAD Imaging Facility for their support in the acquisition of microscopy data.

## DISCLOSURE

All the authors declared no competing interests.

## Supplementary Information

**Supplementary Figure S1.**
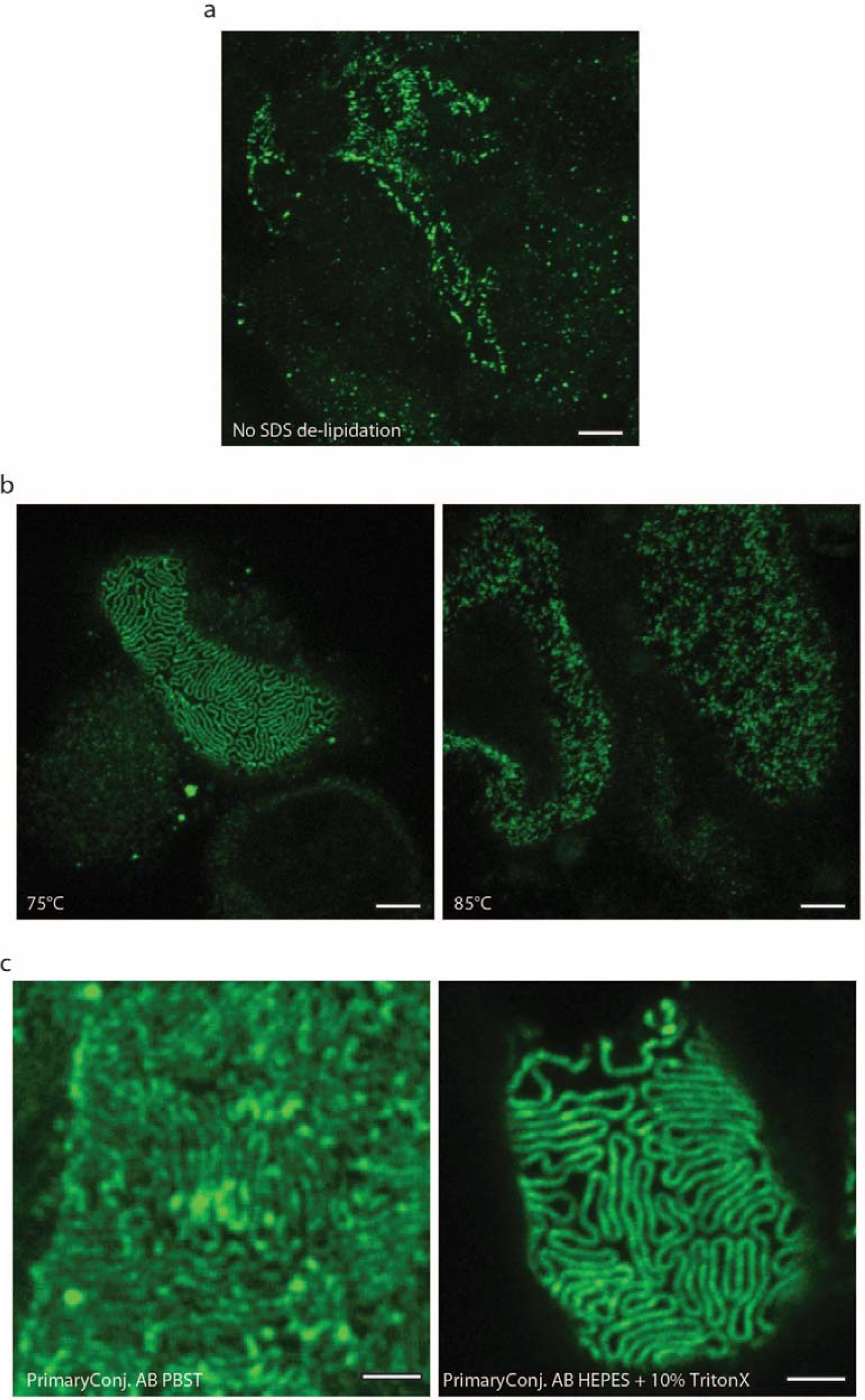
Effect of SDS clearing on labelling quality and sample intergrity. Scale bars 2 µm (a, b), 1 µm (c). (a) Effect of omitting SDS clearing. The sample was treated according to the protocol shown in Fig. 1, but the SDS clearing step was omitted. The sample was stained for nephrin and a z-stack was acquired, which was then maximum intensity projected using Fiji. Due to limited penetration of the antibody and poor nephrin staining, foot processes could not be visualized. (b) Clearing at 85°C alters nanoscale tissue integrity. (c) Effect of diluting primary conjugated nephrin antibodies in HEPES buffer with 10% TritonX as compared to using a standard PBST buffer.

**Supplementary Figure S2.**
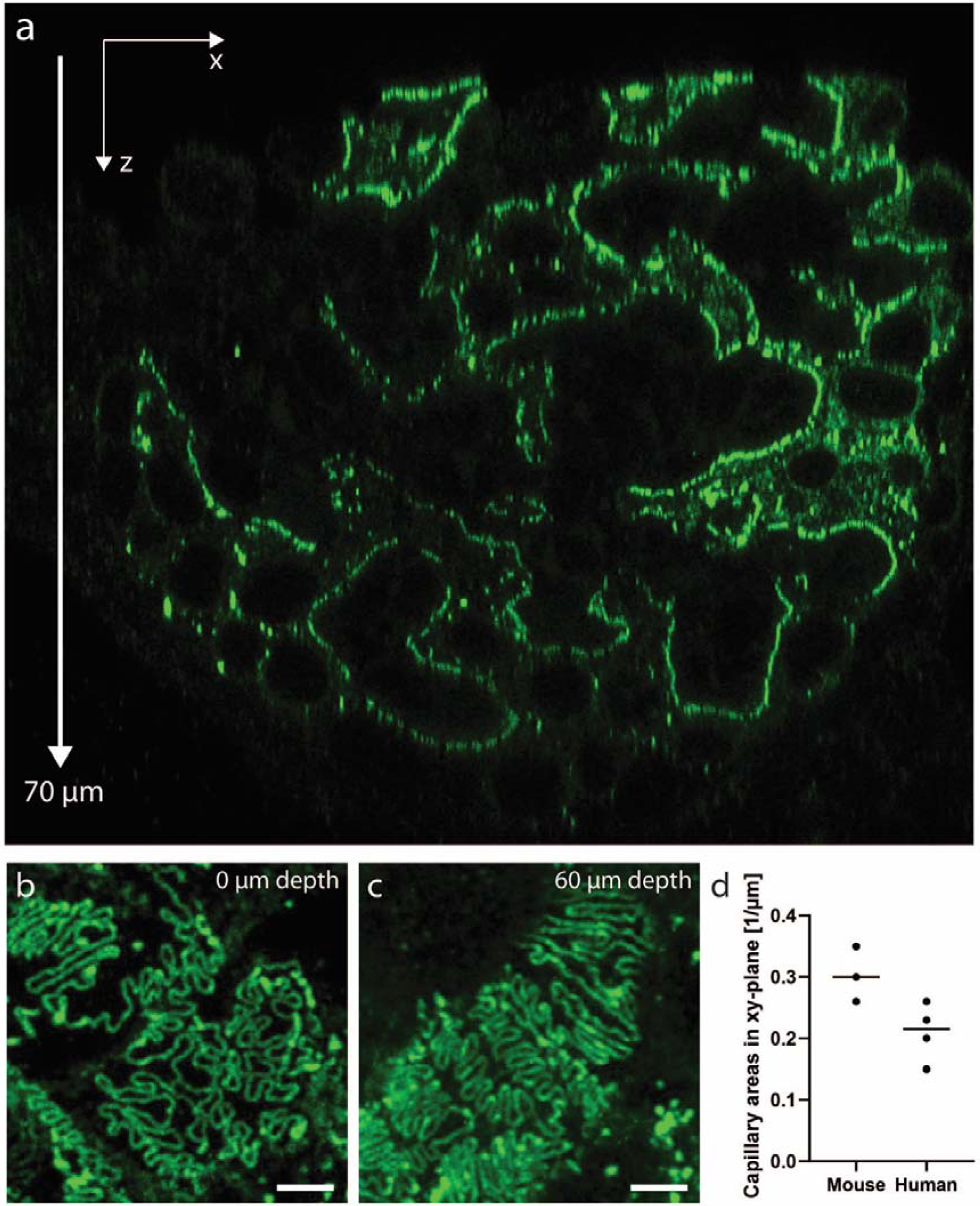
Imaging depth in samples treated according to our protocol. A human kidney sample (nephrectomy due to renal tumor) was stained for nephrin using Alexa-488 and imaged using a Leica SP8 confocal system equipped with a 93X 1.3 NA glycerol objective with a motorized correction collar to correct for spherical aberrations. (a) xz-slice showing that the penetration of the antibodies allows for imaging foot process at least 70 µm into the sample. (b-c) Maximum intensity projections of 2-5 µm confocal z-stacks taken at surface (b) and 60 µm into the sample (c), showing that foot processes outlined by the nephrin staining can can clearly be resolved deep inside human tissue. (d) # of areas per µm in depth where glomerular capillaries are oriented in the xy-plane. Three z-stacks of whole mouse glomeruli, and 4 z-stacks from whole glomeruli in human patients were analyzed.

**Supplementary Figure S3.**
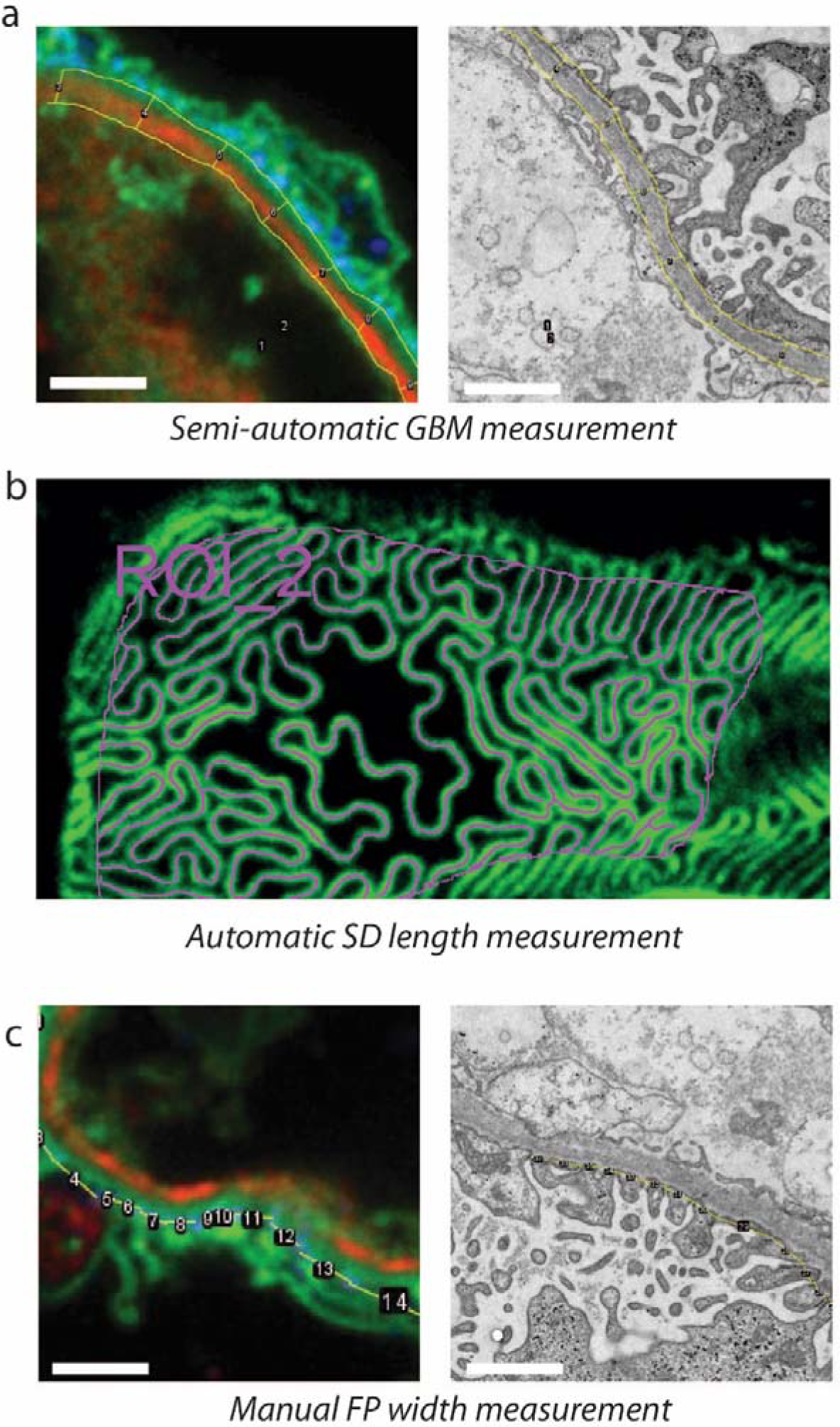
Visual description of quantification methods used in the manuscript. (a) For GBM thickness measurements, an ImageJ macro lets the user outline inner and outer boundaries of the glomerular basement membrane (GBM) and measures the GBM thickness in increments of 1 µm. The inner boundary was taken as the interface between the WGA signal (endothelium) and the collagenIV signal and the outer boundary right under the nephrin signal as shown. (b) For slit diaphragm length measurements, and ImageJ macro lets the user define a region of interest on the glomerular capillary and automatically detects and measures the slit diaphragm length within the area of the ROI. (c) Manual measurement of foot process width in cross-sectional view of glomerular capillaries. The distance between nephrin maxima were taken as the foot process width. No correction for cutting or optical sectioning artifacts were applied since measurements were carried out only for comparative purposes.

**Supplementary Figure S4.**
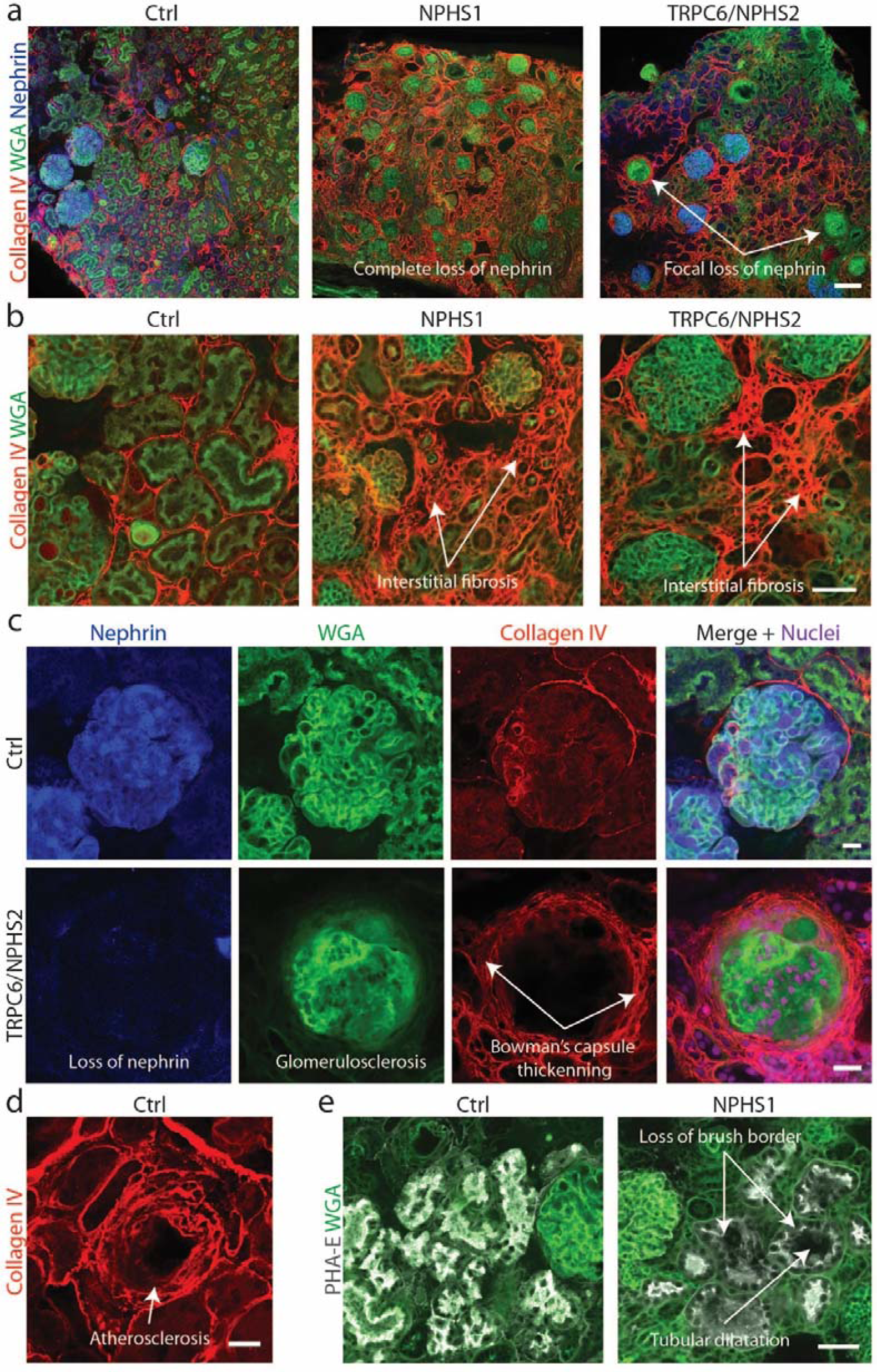
Fluorescence-based large-scale pathology of human samples. (a-e) Confocal images acquired using a Zeiss LSM 780 confocal microscope equipped with a 10X/0.45NA air objective. All samples were stained for nuclei with DRAQ-5, nephrin with Alexa-405, WGA conjugated to Alexa-488, collagen IV with Alexa-555 and PHA-E conjugated to Atto-594. Relevant channels are shown to visualize desired features in each image. (a) Images of control (renal carcinoma) and congenital nephrotic syndrome (NPHS1 and TRPC6/NPHS2 mutations) samples showing overall alterations in morphology and in the expression of nephrin. Scale bar 100 µm. (b) Zoomed images showing different degrees of interstitial fibrosis visualized by collagen IV staining. Scale bar 50 µm. (c) Zoomed images of a healthy glomerulus (nephrectomy patient) and a sclerotic (CNS, TRPC6/NPHS2) glomerulus. Different features of glomerulosclerosis are apparent. Scale bars 20 µm. (d) Zoomed image from the patient that underwent nephrectomy due to renal carcinoma shows clear atherosclerosis visualized by collagen IV staining. Scale bar 20 µm. (e) Zoomed images of tubules in control and NPHS1 CNS patient shows loss of brush border visualized by PHA-E staining and dilatation of tubules and tubular lumen. Scale bar 50 µm.

**Supplementary Figure S5.**
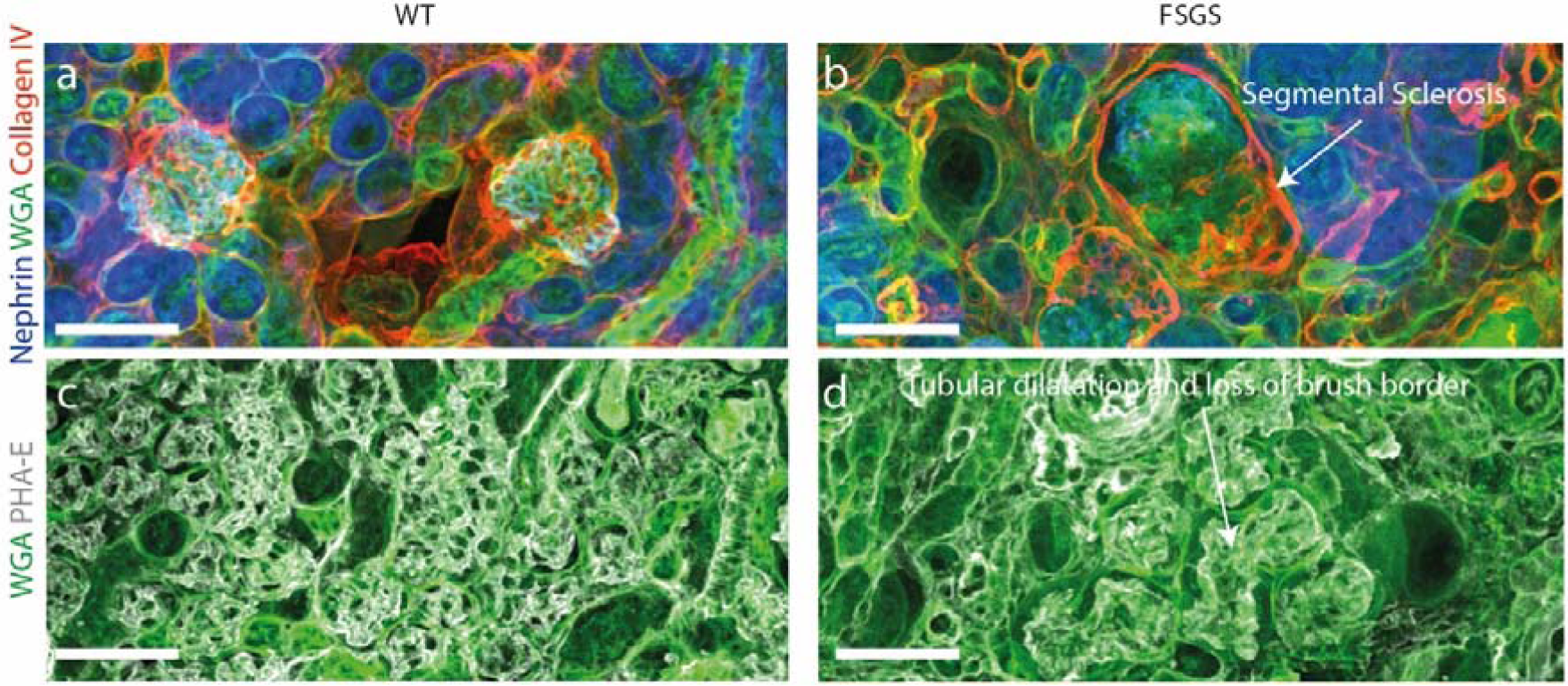
Histological features for the FSGS mouse line. Samples were stained with CF-405M conjugated WGA lectin, an Alexa-488 conjugated nephrin antibody, an Alexa-555 conjugated collagen IV antibody, and Atto594 conjugated PHA-E lectin. Scale bars 100 µm. (a-b) Segmental sclerosis can clearly be visible as lack of nephrin in lower part of the glomerulus in (b). (c-d) Dilatation of tubules as well as a decrease in amount of brush border is apparent from the PHA-E signal (grey).

**Supplementary Table S1.** List of pathology features that can be detected and in some cases quantified using our protocol.

| Pathology feature | Can detect? |
| --- | --- |
| Complete glomerulosclerosis | Yes |
| Segmental glomerulosclerosis | Yes |
| Tubular atrophy | Yes |
| Tubular dilatation | Yes |
| Interstitial Fibrosis | Yes |
| Interstitial infiltration of cells | Yes |
| Distinguishing cell types | Not tested |
| Atherosclerosis | Yes |
| Endothelial hypercellularity | Yes |
| Mesangial expansion | Yes |
| Mesangial hypercellularity | Yes |
| Foot process effacement | Yes |
| GBM thickness | Yes |
| Endothelial alterations | Not tested |
| Fibrillary deposits | No |
| IgA deposits | Yes |
| Other immunoglobulin or complement factors | Not tested |
| Amyloid | Not tested |
| Hyalinosis | Not tested |
| Tubuloreticular inclusions | Not tested |

## Notes

### Competing Interest Statement

The authors have declared no competing interest.

## REFERENCES

1 Chozinski TJ, Mao C, Halpern AR, et al. Volumetric, Nanoscale Optical Imaging of Mouse and Human Kidney via Expansion Microscopy. Sci Rep 2018;8:10396.

2 Pullman JM, Nylk J, Campbell EC, et al. Visualization of podocyte substructure with structured illumination microscopy (SIM): a new approach to nephrotic disease. Biomed Opt Express 2016;7:302–311.

3 Siegerist F, Ribback S, Dombrowski F, et al. Structured illumination microscopy and automatized image processing as a rapid diagnostic tool for podocyte effacement. Sci Rep 2017;7:11473.

4 Suleiman H, Zhang L, Roth R, et al. Nanoscale protein architecture of the kidney glomerular basement membrane. Elife 2013;2. doi:10.7554/eLife.01149.

5 Unnersjö-Jess D, Scott L, Blom H, et al. Super-resolution stimulated emission depletion imaging of slit diaphragm proteins in optically cleared kidney tissue. Kidney Int 2016;89:243–247.

6 Unnersjö-Jess D, Scott L, Sevilla SZ, et al. Confocal super-resolution imaging of the glomerular filtration barrier enabled by tissue expansion. Kidney Int 2018;93:1008– 1013.

7 Zhao Y, Bucur O, Irshad H, et al. Nanoscale imaging of clinical specimens using pathology-optimized expansion microscopy. Nat Biotechnol 2017;35:757.

8 Bucur O, Fu F, Calderon M, et al. Nanoscale imaging of clinical specimens using conventional and rapid-expansion pathology. Nat Protoc 2020;15:1649–1672.

9 Pullman JM. New Views of the Glomerulus: Advanced Microscopy for Advanced Diagnosis. Front Med 2019;6:37.

10 Siegerist F, Endlich K, Endlich N. Novel Microscopic Techniques for Podocyte Research. Front Endocrinol (Lausanne) 2018;9:379.

11 Butt L, Unnersjö-Jess D, Höhne M, et al. A molecular mechanism explaining albuminuria in kidney disease. Nat Metab 2020;2:461–474.

12 Motrapu M, Swiderska MK, Mesas I, et al. Drug Testing for Residual Progression of Diabetic Kidney Disease in Mice Beyond Therapy with Metformin, Ramipril, and Empagliflozin. J Am Soc Nephrol 2020;:ASN.2019070703.

13 Artelt N, Siegerist F, Ritter AM, et al. Comparative Analysis of Podocyte Foot Process Morphology in Three Species by 3D Super-Resolution Microscopy. Front Med 2018;5:292.

14 Moeller MJ, Chia-Gil A. A step forward in understanding glomerular filtration. Nat Rev Nephrol 2020. doi:10.1038/s41581-020-0313-6.

15 Xu N, Tamadon A, Liu Y, et al. Fast free-of-acrylamide clearing tissue (FACT)—an optimized new protocol for rapid, high-resolution imaging of three-dimensional brain tissue. Sci Rep 2017;7:9895.

16 Murakami TC, Mano T, Saikawa S, et al. A three-dimensional single-cell-resolution whole-brain atlas using CUBIC-X expansion microscopy and tissue clearing. Nat Neurosci 2018;21:625–637.

17 Hou B, Zhang D, Zhao S, et al. Scalable and DiI-compatible optical clearance of the mammalian brain. Front Neuroanat 2015;9:19.

18 Susaki EA, Shimizu C, Kuno A, et al. Versatile whole-organ/body staining and imaging based on electrolyte-gel properties of biological tissues. Nat Commun 2020;11:1982.

19 Wilson T. Resolution and optical sectioning in the confocal microscope. J Microsc 2011;244:113–121.

20 Tryggvason K. Unraveling the Mechanisms of Glomerular Ultrafiltration. Jasn 1999;10:2440–2445.

21 Ramage IJ, Howatson AG, Mccoll JH, et al. Glomerular basement membrane thickness in children: A stereologic assessment. Kidney Int 2002;62:895–900.

22 Amann K, Haas CS. What you should know about the work-up of a renal biopsy. Nephrol Dial Transplant 2006;21:1157–1161.

23 Sachs W, Sachs M, Krüger E, et al. Distinct Modes of Balancing Glomerular Cell Proteostasis in Mucolipidosis Type II and III Prevent Proteinuria. J Am Soc Nephrol 2020;:ASN.2019090960.

24 Suleiman HY, Roth R, Jain S, et al. Injury-induced actin cytoskeleton reorganization in podocytes revealed by super-resolution microscopy. JCI insight 2017;2:e94137.

25 Nyström S, Bäck M, Nilsson KPR, et al. Imaging amyloid tissues stained with luminescent conjugated oligothiophenes by hyperspectral confocal microscopy and fluorescence lifetime imaging. J Vis Exp 2017;2017:e56279.

26 Torres R, Velazquez H, Chang JJ, et al. Three-Dimensional Morphology by Multiphoton Microscopy with Clearing in a Model of Cisplatin-Induced CKD. J Am Soc Nephrol 2016;27:1102.

27 Puelles VG, Fleck D, Ortz L, et al. Novel 3D analysis using optical tissue clearing documents the evolution of murine rapidly progressive glomerulonephritis. Kidney Int 2019. doi:10.1016/J.KINT.2019.02.034.

28 Rivenson Y, Wang H, Wei Z, et al. Virtual histological staining of unlabelled tissue-autofluorescence images via deep learning. Nat Biomed Eng 2019;3:466–477.

29 Hermsen M, Bel T, Boer M Den, et al. Deep learning-based histopathologic assessment of kidney tissue. J Am Soc Nephrol 2019;30:1968–1979.

30 Ginley B, Lutnick B, Jen KY, et al. Computational segmentation and classification of diabetic glomerulosclerosis. J Am Soc Nephrol 2019;30:1953–1967.

31 Graham L, Orenstein JM. Processing tissue and cells for transmission electron microscopy in diagnostic pathology and research. Nat Protoc 2007;2:2439.

